# A conformational sensor based on genetic code expansion reveals an autocatalytic component in EGFR activation

**DOI:** 10.1101/314682

**Authors:** Martin Baumdick, Márton Gelléri, Chayasith Uttamapinant, Václav Beránek, Jason W Chin, Philippe IH Bastiaens

## Abstract

Epidermal growth factor receptor (EGFR) activation by growth factors (GFs) relies on dimerization and allosteric activation of its intrinsic kinase activity, resulting in trans-phosphorylation of tyrosines on its C-terminal tail. While structural and biochemical studies identified this EGF-induced allosteric activation, imaging collective EGFR activation in cells and molecular dynamics simulations pointed at additional catalytic EGFR activation mechanisms. To gain more insight in EGFR activation mechanisms in living cells, we developed a Förster Resonance Energy Transfer (FRET) based <u>con</u>formational <u>EG</u>FR <u>i</u>ndicator (CONEGI) using genetic code expansion that reports on conformational transitions in the EGFR activation loop. Comparing conformational transitions, self-association and auto-phosphorylation of CONEGI and its Y845F mutant revealed that Y_845_ phosphorylation induces a catalytically active conformation in EGFR monomers. This conformational transition depends on EGFR kinase activity and auto-phosphorylation on its C–terminal tail, generating a looped causality that leads to autocatalytic amplification of EGFR phosphorylation at low EGF dose.

---

Dimerization of EGFR by GFs activates its intrinsic kinase activity, which trans-phosphorylates tyrosine residues on the C-terminal receptor tail^1,2^. SH2- or PTB-containing signal transducing proteins are then recruited to these phosphorylated tyrosines, propagating the signal in the cytoplasm^3,4^. Structural data of EGFR indicate that in absence of ligand a closed tethered extracellular domain (ECD) and association of the intracellular tyrosine kinase domain (TKD) with the negatively charged plasma membrane (PM) by two polybasic stretches favor steric auto-inhibition of EGFR’s intrinsic kinase activity^5-7^. Ligand binding to EGFR is coupled to conformational changes in the extra- and intracellular domains and overcomes intrinsic auto-inhibition resulting in allosteric activation via asymmetric dimer formation of the TKD^2,5^. For this, the αC-helix located in the N lobe of the TKD moves from its ‘out’-configuration to an ordered ‘in’-configuration, while the activation loop frees the catalytic cleft and undergoes conformational rearrangements of ~20 Å^8-10^. Despite the steric auto-inhibitory features, autonomous EGFR phosphorylation was observed in several cancer types including breast and lung cancer that either exhibit high EGFR surface concentrations through EGFR overexpression or bear oncogenic mutations favoring an active conformation^9, 11-13^. Spontaneous auto-phosphorylation of unliganded EGFR can occur due to thermal fluctuations that overcome intrinsic steric auto-inhibition^14-16^. These auto-phosphorylation events can trigger an autocatalytic amplification mechanism when they induce an active conformation that further catalyzes EGFR auto-phosphorylation^15^. Molecular dynamics simulations suggested that Y_845_ phosphorylation in the EGFR activation loop suppresses intrinsic disorder in the αC-helix thereby stabilizing an active receptor conformation as well as increasing EGFR dimerization^9^. We therefore investigated whether EGFR can adopt an active conformation upon Y_845_ phosphorylation and how this impacts on collective EGFR phosphorylation dynamics in living cells. A clear avenue to obtain a better insight in collective EGFR activation is to monitor conformational dynamics of the TKD. For this, we engineered a FRET-based <u>con</u>formational <u>EG</u>FR <u>i</u>ndicator (CONEGI) using genetic code expansion. In contrast to existing kinase activity sensors based on substrate phosphorylation^17^, CONEGI was designed to report on conformational transitions in a functional domain of the EGFR TKD by the change in distance and orientation of a fluorophore conjugated to an unnatural amino acid (UAA) relative to the fluorescent protein mCitrine inserted into a conformationally invariant region. Based on structural data, we identified the end of the TKD as an insertion site for mCitrine that is conformationally invariant and does not affect EGFR function. UAA incorporation and subsequent site specific labeling at position 851 created a FRET-based sensor that reports on conformational transitions of the EGFR activation loop. This construct retained EGFR dimerizing and catalytic functionality. Monitoring conformational transitions together with dimerization and auto-phosphorylation and comparing these readouts to a CONEGI Y845F mutant revealed that an active conformation in monomeric receptors is induced by Y_845_ phosphorylation. We then show that Y_845_ phosphorylation depends on auto-phosphorylation of the C-terminal tail, which creates an autocatalytic loop that amplifies EGFR phosphorylation at low, non-saturating EGF concentrations.

## Results

### Design and performance of CONEGI

To monitor conformational states of the TKD we engineered multiple FRET-based conformational EGFR sensor variants, in which the donor, monomeric Citrine (mCitrine), was always genetically encoded at the same rigid region of the TKD and genetic code expansion and bioorthogonal labeling chemistry were used to position the acceptor, Atto590, at different, flexible structures of the TKD that change conformation upon activation (**Figure 1a**). This hybrid sensor design allows measuring structural changes in different key functional TKD regions relative to the donor. Conformational movements of these TKD regions will alter the distance and angle between mCitrine and Atto590 resulting in changes in FRET efficiency, which can be quantified by fluorescence lifetime imaging microscopy (FLIM). mCitrine was selected as donor because of its mono-exponential fluorescence decay profile^18,19^ and the membrane-permeable Atto590 as acceptor because of its high extinction coefficient (ɛ: 120000), high quantum yield (QY: 0.8) and spectral overlap with mCitrine (R_0_=5.9 nm).

**Figure 1:**
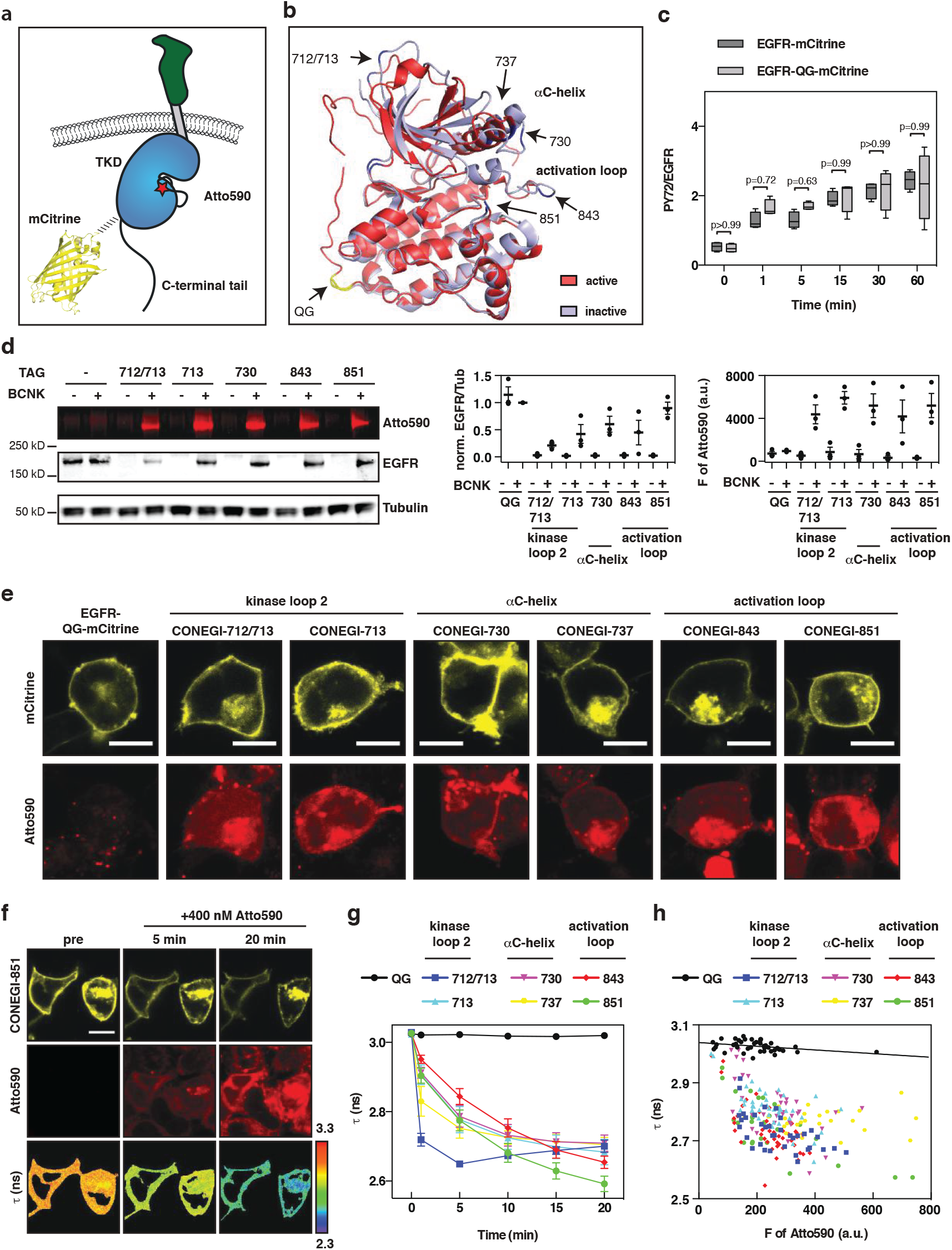
Design and performance of CONEGI. (**a**) Schematic representation of CONEGI. mCitrine is fused to the C-terminal end of the tyrosine kinase domain (TKD) using a coiled-coil linker (dashed line) and Atto590 (red star) is site-specifically attached to the activation loop. (**b**) Alignment of active (red; PDB: 2J5F) and inactive (cyan; PDB: 2GS7) crystal structures of the EGFR TKD. mCitrine insertion (QG, yellow) and BCNK incorporation sites (blue, black arrows) are indicated. (**c**) Relative phosphorylation (PY72/EGFR) of EGFR-mCitrine or EGFR-QG-mCitrine upon EGF stimulation determined by Western blot analysis (n=4) (**Supplementary Figure 1b**). (**d**) Fluorescence images and Western blot analysis following SDS-PAGE of HEK293T cell lysates showing Atto590 fluorescence and expression level of EGFR-QG-mCitrine and CONEGIs depending on BCNK. Blots were probed with anti-EGFR and anti-Tubulin (left). Normalized relative EGFR expression (EGFR/Tub) (middle) and Atto590 fluorescence intensity (right) of EGFR-QG-mCitrine and CONEGIs (n=3 blots). (**e**) Representative mCitrine and Atto590 fluorescence images of EGFR-QG-mCitrine and CONEGIs in HEK293T cells. (**f**) Representative fluorescence images of EGFR(BCNK851)-QG-mCitrine upon tetrazine-Atto590 labeling and corresponding τ. (**g**) Mean τ in CONEGIs and EGFR-QG-mCitrine (QG: n=10 cells; 712/713: n=8; 713: n=6; 730: n=11; 737: n=7; 843: n=10; 851: n=17) at the PM upon tetrazine-Atto590 addition. (**h**) Dependency of τ in CONEGIs and EGFR-QG-mCitrine (QG: n=42 cells; 712/713: n=36; 713: n=43; 730: n=36; 737: n=30; 843: n=44; 851: n=32) on mean Atto590 fluorescence intensity (F of Atto590) at the PM after 20 min labeling. Data points represent individual cells. τ, fluorescence lifetime of mCitrine. Scale bars, 10 μm. EGF stimulation, 100 ng/ml. Error bars: standard error of the mean, except **Figure 1c**: tukey box plot.

By aligning active (red; PDB: 2J5F) and inactive (cyan; PDB: 2GS7) crystal structures of the EGFR TKD (**Figure 1b**), we identified a conformationally invariant region at the TKD end, where we inserted mCitrine between amino acids (aa) Q958 and G959 (EGFR-QG-mCitrine)^15^. This site is exposed to the protein surface and not part of the dimerization interface or regions known to be essential for kinase activity (**Supplementary Figure 1a**). To further minimize perturbation of the TKD structure and constrain the mCitrine orientation, we used two linkers that form an antiparallel coiled coil helix for mCitrine insertion^20^. EGFR-QG-mCitrine was fully active and correctly localized as apparent from its similar localization and EGF-induced phosphorylation as compared to C-terminally tagged EGFR (EGFR-mCitrine) (**Figure 1c; Supplementary Figure 1b,c**), which was shown to follow the localization and activity of endogenous EGFR^21,22^. We then selected three regions (kinase loop 2, αC-helix and activation loop) that showed substantial differences between the active and inactive conformation for Atto590 attachment (**Figure 1b**). We replaced the coding sequences of amino acids K713 in the kinase loop 2, K730 and D737 in the αC-helix and K843 and K851 in the activation loop with an amber codon and also inserted an amber codon between the coding sequence for E712 and K713. This resulted in the creation of a series of EGFR(TAGXXX)-QG-mCitrine variants, where XXX indicates the amber codon position. We estimated the distances between the QG site that included the maximum linker length and each proposed site of Atto590 attachment. The obtained distances (R) of ~7.4-8.5 nm were sufficiently close to the Förster radius (R_0_=5.9 nm) of the employed FRET pair resulting in estimated FRET efficiencies (E_FRET_) between 10-21% for the different variants as calculated by 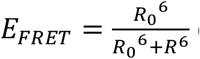 (**Supplementary Table 1**). This showed that the 851-site yielded the highest (21%) FRET efficiency. We conjugated Atto590 to the desired site in the TKD in a two-step process to generate conformational EGFR indicators (CONEGIs): first, we used a derivative of the pyrrolylsyl-tRNA synthetase/tRNA_CUA_ pair from *Methanosarcina mazei* to express each EGFR(TAGXXX)-QG-mCitrine gene and direct the incorporation of added unnatural amino acid Bicyclo[6.1.0]nonyne-lysine (BCNK)^23,24^, second, we labeled each EGFR(BCNKXXX)-QG-mCitrine variant, where XXX indicates the position of BCNK incorporation, with a tetrazine-Atto590 (tet-Atto590) conjugate via an inverse electron demand Diels Alder reaction.

Expression of EGFR(BCNKXXX)-QG-mCitrine variants in HEK293T cells was dependent on BCNK (**Figure 1d**). Whereas EGFR(BCNK851)-QG-mCitrine was expressed at a comparable level to EGFR-QG-mCitrine, all other variants exhibited reduced expression (**Figure 1d**). EGFR(BCNKXXX)-QG-mCitrine labeling with tet-Atto590 was selective, as judged by both, fluorescence imaging of cell lysates following SDS PAGE, and co-localization of mCitrine and Atto590 fluorescence at the PM and intracellular compartments as visualized in living cells by confocal microscopy (**Figure 1d,e**). All EGFR(BCNKXXX)-QG-mCitrine variants exhibited a similar localization as compared to an EGFR variant C-terminally tagged with mTurquoise (EGFR-mTurquoise) (**Supplementary Figure 1d**). In addition to their comparable PM distribution (**Supplementary Figure 1d,e**), all CONEGI constructs exhibited a pericentriolar localization similar to EGFR-mTurquoise. This pericentriolar compartment was identified to be the Rab11-positive recycling endosome (**Supplementary Figure 1f**), which was previously shown to maintain EGFR at the PM by continuous recycling^15,25^.

To experimentally determine whether EGFR(BCNKXXX)-QG-mCitrine labeling with Atto-590 results in FRET, we imaged the fluorescence lifetime (τ) of mCitrine after addition of tet-Atto590 to living cells by FLIM. For all CONEGI’s we obtained a significant decrease in τ of mCitrine from 3.02 ± 0.004 ns to 2.67 ± 0.039 ns over a 20 min time course, corresponding to an average E_FRET_ between 10-14% as calculated by: 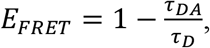 where Atto590 conjugation to BCNK incorporated into the 851 site yielded the highest FRET efficiency (14.3 ± 0.8%) (**Figure 1f,g**). These experimental E_FRET_ values were in broad agreement with the theoretical predictions, exhibiting equal or lower values possibly due to the relative orientation between the dyes, which affects the parameter κ^2^ in R_0_ (**Supplementary Table 1, Figure 1g**). CONEGI-712/713 exhibited the fastest labeling kinetics, with a reaction time clearly below 5 min, which likely reflects the high accessibility of this site for fluorophore attachment (**Figure 1g**). The drop in τ of mCitrine associated with tet-Atto590 conjugation was reversed for all CONEGI’s upon Atto590 photobleaching (**Supplementary Figure 1g,h**). Furthermore, tet-Atto590 addition to cells co-expressing EGFR-QG-mCitrine and the BCNK incorporation system in presence of BCNK did not lead to an alteration in τ of mCitrine (**Figure 1g,h**). This demonstrated that the obtained changes in τ in the CONEGI’s result from specific FRET between mCitrine and Atto590 conjugated to site-specifically incorporated BCNK and not from tet-Atto590 non-specifically bound to EGFR or the PM, or from labeling of BCNK that might be incorporated at genomic amber codons. Washout of unbound tet-Atto590 after labeling of cells did not abolish the decrease in τ of mCitrine for the CONEGI’s, confirming that the conjugation of tet-Atto590 to BCNK is stable (**Supplementary Figure 1i**). The negative correlation of τ with the Atto590 fluorescence intensity and the saturation of the binding curves at high Atto590 concentrations for all CONEGI’s further confirmed specificity of tet-Atto590-conjugation to BCNK in EGFR (**Figure 1h**).

### FRET changes report on conformational transitions of the activation loop in CONEGI-851

To investigate whether the CONEGI’s report on conformational changes in the TKD that occur upon activation, we measured τ of mCitrine at the PM by FLIM following stimulation with EGF to cells. We observed a significant decrease in τ of CONEGI-737, −843 and −851 and a significant increase in τ of CONEGI-712/713 upon EGF stimulation, whereas τ of CONEGI-713 and −730 did not significantly change (**Figure 2a; Supplementary Figure 2a**). To account for the variability in the completeness of the Atto590 labeling reaction that affects the initial FRET efficiency in a particular CONEGI construct, the difference in τ (Δτ) relative to the τ before stimulation in each experiment was plotted (**Figure 2b**). This Δτ-time plot that follows the general trends of the τ-time plot (**Supplementary Figure 2a**) clearly reflects the conformational transitions of the CONEGI constructs. Neither τ of EGFR-QG-mCitrine in presence of tet-Atto590 nor τ of EGFR(BCNK851)-QG-mCitrine in absence of tet-Atto590 did change upon EGF addition, precluding photophysical effects that change τ of mCitrine upon EGF stimulation (**Figure 2b**; **Supplementary Figure 2b**).

**Figure 2:**
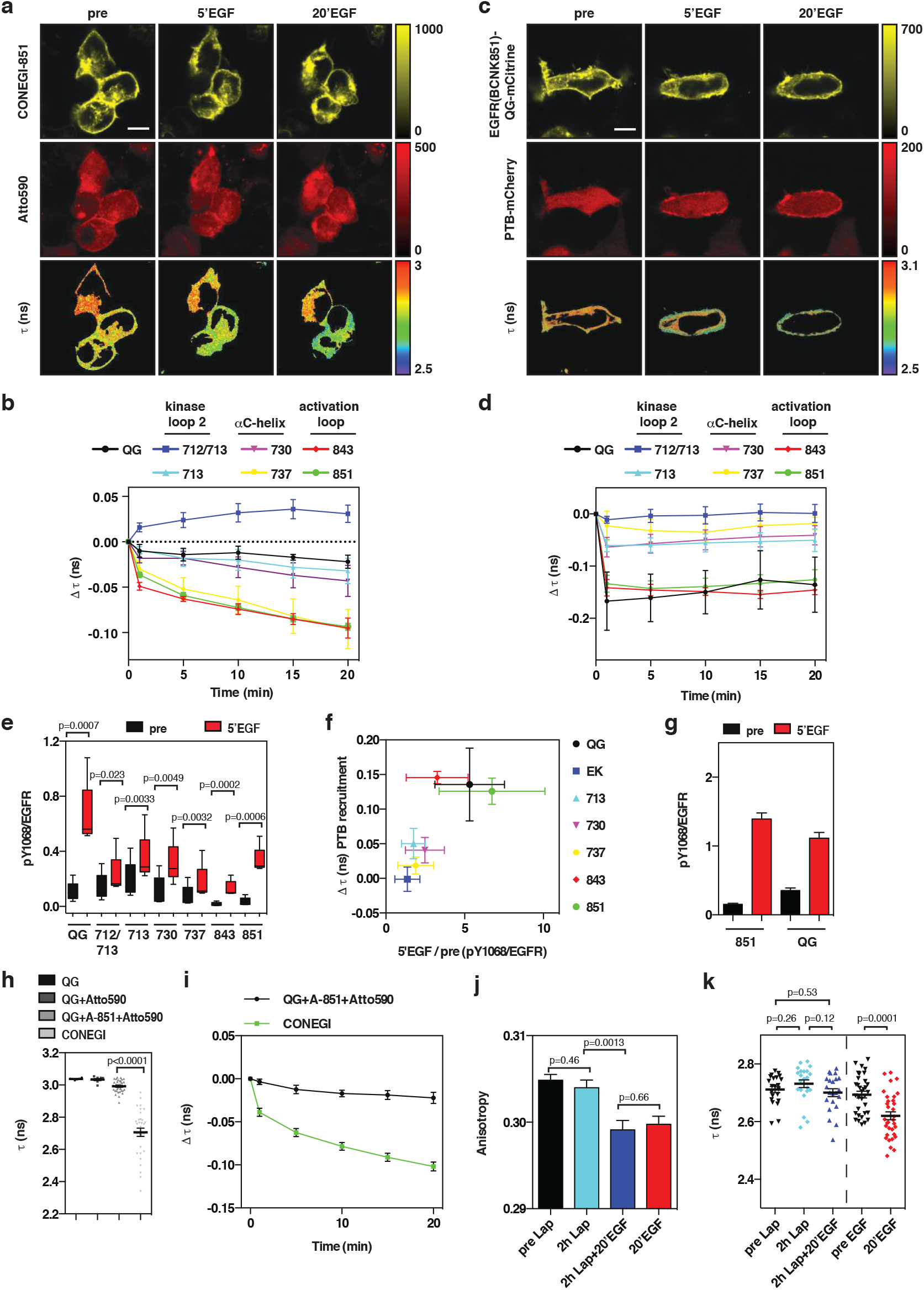
CONEGI-851 reports on conformational changes in the activation loop. (**a**) Representative CONEGI-851 fluorescence and corresponding τ images upon EGF stimulation. (**b**) Change in Δτ of CONEGIs and EGFR-QG-mCitrine (QG: n=5 cells; 712/713: n=7; 713: n=7; 730: n=7; 737: n=7; 843: n=6; 851: n=31) at the PM upon EGF stimulation (**Supplementary Figure 2a**: absolute τ). (**c**) Representative fluorescence images of EGFR(BCNK851)-QG-mCitrine, PTB-mCherry and corresponding τ upon EGF stimulation. (**d**) Change in Δτ of EGFR-QG-mCitrine or EGFR(BCNKXXX)-QG-mCitrine variants (QG: n=4 cells; 712/713: n=6; 713: n=8; 730: n=8; 737: n=5; 843: n=6; 851: n=6) at the PM upon EGF-mediated PTB-mCherry recruitment. (**e**) Relative Y_1068_ phosphorylation (pY_1068_/EGFR) of EGFR-QG-mCitrine and CONEGIs upon EGF stimulation by Western blot analysis (n=5) (**Supplementary Figure 2c**). (**f**) Fold-change in Y_1068_ phosphorylation (5’EGF/pre) (quantified from **e**) versus Δτ upon PTB-mCherry recruitment (quantified from **d**) for CONEGIs and EGFR-QG-mCitrine. (**g**) Relative Y_1068_ phosphorylation (pY_1068_/EGFR) of EGFR-QG-mCitrine (QG) and EGFR(BCNK851)-QG-mCitrine (851) at the PM upon EGF stimulation by immunofluorescence (851: pre: n=72 cells; 5’ÉGF: n=67: QG: pre: n=73; 5’ÉGF: n=74) (**Supplementary Figure 2d**). (**h**) τ of EGFR-QG-mCitrine in absence (QG; n=6 cells) or presence of tetrazine-Atto590 (QG+Atto590; n=9), EGFR-QG-mCitrine co-expressed with Atto590-labeled EGFR(BCNK851) (QG+A-851+Atto590; n=36) and CONEGI (n=32) (**Supplementary Figure 2f**). (**i**) Change in Δτ of CONEGI (n=27 cells) or EGFR-QG-mCitrine co-expressed with Atto590-labeled EGFR(BCNK851) (n=7) upon EGF stimulation. (**j**) Mean mCitrine fluorescence anisotropy of CONEGI upon EGF stimulation in presence or absence of 1 μM Lapatinib (Lap) (**Supplementary Figure 2h**) (N=3 experiments). (**k**) τ of CONEGI at the PM upon EGF stimulation in presence (left; n=22 cells) or absence (right; n=33) of 1 μM Lap. Scale bars, 10 μm. EGF stimulation, 100 ng/ml. Error bars: standard error of the mean, except **Figure 2e**: tukey box plot. τ, fluorescence lifetime of mCitrine.

To examine whether the EGFR(BCNKXXX)-QG-mCitrine variants retain their activity upon BCNK incorporation, we quantified the phosphorylation on tyrosines 1086 (Y_1086_) and 1148 (Y_1148_) by measuring the interaction of mCherry-tagged phosphotyrosine-binding domain (PTB-mCherry) with each EGFR(BCNK)-QG-mCitrine variant upon EGF stimulation by FLIM^26^. EGF-induced PTB-mCherry recruitment to EGFR(BCNK843)-QG-mCitrine and EGFR(BCNK851)-QG-mCitrine at the PM resulted in a corresponding drop in τ (**Figure 2c,d**). This decrease in τ was comparable to that of EGFR-QG-mCitrine upon EGF-mediated PTB-mCherry recruitment, whereas EGF-induced phosphorylation of other variants (712/713, 713, 730, and 737) only marginally increased as judged by minimal decreases in τ of mCitrine (**Figure 2d**). To examine whether other phosphorylation sites were affected by BCNK incorporation and Atto590-labeling, we quantified the relative phosphorylation (pY/EGFR) on the Grb2-binding site Y_1068_^27^ for all CONEGIs by Western blot analysis. Consistent with PTB-mCherry recruitment, CONEGI-843 and −851 exhibited a similar EGF-induced fold-change in Y_1068_ phosphorylation as compared to EGFR-QG-mCitrine, but the relative phosphorylation level of CONEGI-843 was drastically reduced (**Figure 2e,f; Supplementary Figure 2c**). The other CONEGIs showed increased autonomous phosphorylation (CONEGI-712/713 and −713) or responded only marginally to EGF (CONEGI-712/713, −713, −730 and −737), indicating that their activation mechanism was impaired (**Figure 2e,f; Supplementary Figure 2c**). Immunofluorescence staining against pY_1068_ in fixed HEK293T cells expressing EGFR(BCNK851)-QG-mCitrine showed that EGF-mediated phosphorylation at the PM was comparable to that of EGFR-QG-mCitrine (**Figure 2g; Supplementary Figure 2d**), which shows that EGFR(BCNK851)-QG-mCitrine at the PM is fully functional. Tet-Atto590-labeling did not affect autonomous or ligand-dependent EGFR phosphorylation of EGFR(BCNK851)-QG-mCitrine (**Supplementary Figure 2e**). We therefore conclude that CONEGI-851 (from now on denoted as CONEGI) most closely follows the native activation mechanism of EGFR.

To investigate whether the change in FRET upon EGFR activation detected in CONEGI is of intra- or intermolecular origin, we measured τ of EGFR-QG-mCitrine (donor only) upon co-expression with an EGFR(BCNK851) variant that was labeled with Atto590 (acceptor only). τ of EGFR-QG-mCitrine in the presence of Atto590-labeled EGFR(BCNK851) (2.983 ± 0.008) was close to that of EGFR-QG-mCitrine (3.021 ± 0.009) and only marginally decreased upon EGF stimulation (**Figure 2h,i; Supplementary Figure 2f**), showing that FRET in CONEGI reports on intramolecular conformational rearrangements. To demonstrate that the change in FRET efficiency in CONEGI originates from the rearrangement of the activation loop rather than from mCitrine reorientation upon dimerization we locked the TKD into an inactive conformation using the ATP analogue EGFR inhibitor Lapatinib^28^. This abolished EGF-induced CONEGI phosphorylation (**Supplementary Figure 2g**) but did not prevent EGF-induced dimerization as measured by homo-FRET between mCitrine using fluorescence anisotropy^29^ (**Figure 2j; Supplementary Figure 2h)**. Importantly, this EGF-induced dimerization of CONEGI with a Lapatinib locked TKD conformation did not result in a decrease in τ as observed for CONEGI in absence of inhibitor (**Figure 2k**). This shows that CONEGI exclusively reports conformational transitions of the activation loop, but not reorientation of mCitrine upon dimerization.

### Y_845_ phosphorylation induces a catalytically competent conformation of CONEGI monomers

Phosphorylation of Y_845_ was previously described to affect both the conformation of the EGFR TKD, and to enhance its dimerization^9^. We therefore explored if Y_845_ phosphorylation in the activation loop is linked to an EGFR conformational state that is catalytically active without dimerization. For this, we compared the dimerization and conformational transition of CONEGI upon its ligand-independent phosphorylation to that of a CONEGI-Y845F mutant. Ligand-independent phosphorylation of EGFR is known to occur upon phosphatase inhibition by pervanadate (PV)^30^ or upon high EGFR expression levels^14,15,31,32^. Under both conditions the intrinsic EGFR kinase activity surpasses the reverse dephosphorylating activity of phosphatases, resulting in an increased steady state phosphorylation level of EGFR^15^.

EGF administration at saturating dose (100 ng/ml) resulted in a drop in anisotropy of CONEGI (independent of UAA incorporation or Atto590 labeling), showing EGF-induced dimerization (**Figure 3a; Supplementary Figure 3a,b**). This led to a concurrent increase in phosphorylation of Y_1068_ and Y_845_ (**Figure 3b,c; Supplementary Figure 3c,d**). In contrast, the anisotropy of CONEGI did not change upon PV treatment over a time course of 20 min, in which Y_1068_ and Y_845_ phosphorylation reached similar levels as compared to EGF stimulation (**Figure 3a-c; Supplementary Figure 3a,c,d**). This demonstrates that the majority of ligand-independent phosphorylated CONEGI is monomeric^15,33^. Both EGF and PV induced a conformational change in CONEGI as apparent from the decrease in τ. However, while EGF provoked a rapid drop in τ, PV treatment resulted in a slower decrease in agreement with the differential phosphorylation kinetics (**Figure 3b-d; Supplementary Figure 3c-f**). This shows that not only EGF-stimulated dimers but also phosphorylated monomers undergo a conformational change in the activation loop upon phosphorylation. For CONEGI-Y845F expressing HEK293T cells a significantly smaller and slower change in τ upon PV was observed showing that Y845 phosphorylation induces this conformational change in CONEGI monomers (**Figure 3f; Supplementary Figure 3g**). This was paralleled by a significantly lower Y_1068_ auto-phosphorylation in CONEGI-Y845F as compared to that of CONEGI, revealing that Y_845_ phosphorylation results in a catalytically more active CONEGI conformation (**Figure 3e; Supplementary Figure 3h**).

**Figure 3:**
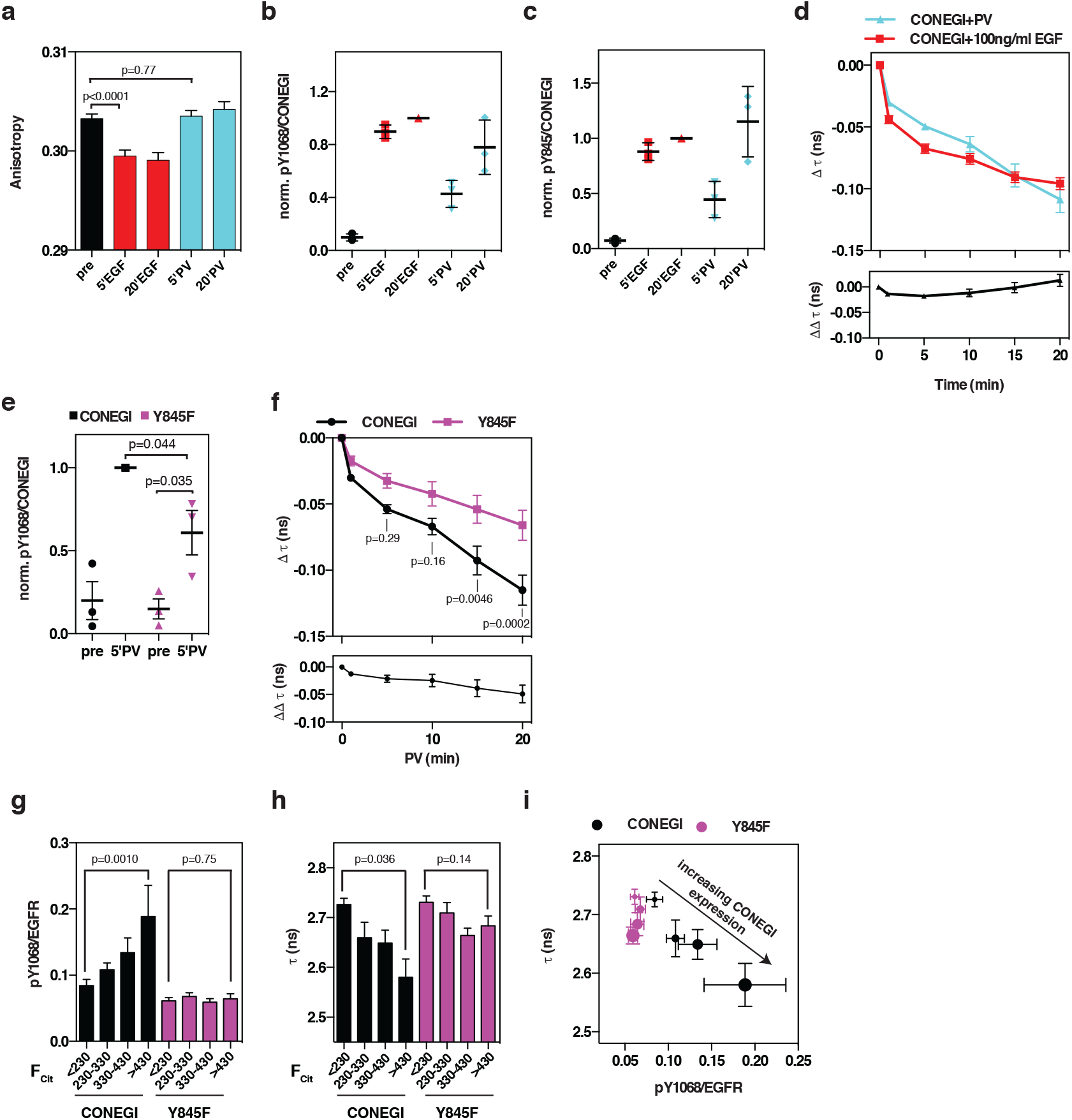
Y_845_ phosphorylation induces an active conformation of EGFR monomers. (**a**) Mean mCitrine fluorescence anisotropy of CONEGI upon EGF or pervanadate (PV) treatment (**Supplementary Figure 3a,b**) (N=3 experiments). (**b,c**) Normalized relative Y_1068_ (pY_1068_/CONEGI) (b) or pY_845_ phosphorylation (pY_845_/CONEGI) (**c**) of CONEGI upon EGF or PV treatment by Western blot analysis (n=3) (**Supplementary Figure 3c,d**). (**d**) Change in Δτ of CONEGI (upper panel) at the PM and the difference between EGF and PV treatment (ΔΔτ) (lower panel) (EGF: n=19 cells; PV: n=6) (**Supplementary Figure 3e,f**). (**e**) Normalized relative Y_1068_ phosphorylation (pY_1068_/CONEGI) of CONEGI or CONEGI-Y845F upon PV treatment by Western blot analysis (n=3) (**Supplementary Figure 3h**). (**f**) Change in Δτ of CONEGI and CONEGI-Y845F upon PV treatment (upper panel) and the difference (ΔΔτ) between CONEGI and CONEGI-Y845F (lower panel) (CONEGI: n=6 cells; Y845F: n=8) (**Supplementary Figure 3g**). (**g**) Relative Y_1068_ phosphorylation (pY_1068_/EGFR) of CONEGI and CONEGI-Y845F at increasing EGFR expression levels as measured by mCitrine intensity per cell (F_cit_ binned as follows: <230, 230-330, 330-430 >430) (CONEGI: n=80 cells; Y845F: n=81). (**h**) Mean τ in CONEGI and CONEGI-Y845F at increasing EGFR expression levels (n=47 cells/variant). (**i**) Relative Y_1068_ phosphorylation (pY_1068_/EGFR) of CONEGI and CONEGI-Y845F versus their τ of mCitrine upon increasing expression levels (F_cit_ <230, 230-330, 330-430 >430; as indicated by increasing dot size). EGF stimulation, 100 ng/ml. PV treatment, 0,33mM. Error bars: standard error of the mean. τ, fluorescence lifetime of mCitrine.

We next investigated the dependence of ligand independent phosphorylation on the expression level of CONEGI in relation to the conformational transition of the activation loop. Exploiting the cell-to-cell variance in mCitrine fluorescence, allowed us to relate CONEGI expression to phosphorylation on Y_1068_ as measured by immunofluorescence. We could clearly observe an expression level dependent increase in auto-phosphorylation that was absent in its Y845F mutant (**Figure 3g,i; Supplementary Figure 3i**). This increase in auto-phosphorylation was not due to an increase in dimers at higher expression as apparent from the fluorescence anisotropy as function of CONEGI expression (**Supplementary Figure 3a**). In an analogous experiment, we related CONEGI expression by its fluorescence intensity to its conformational state by fluorescence lifetime in many individual cells. Here, we could clearly observe a CONEGI expression level dependent conformational transition that was absent in the CONEGI-Y845F mutant (**Figure 3h,i**). Together, these data show that CONEGI but not its Y845F mutant is able to activate other CONEGI molecules via Y_845_ phosphorylation, resulting in catalytic amplification.

### EGFR dimers can induce autocatalytic activation of EGFR monomers

We next investigated whether ligand induced EGFR dimers can activate receptor monomers. At saturating EGF dose (100 ng/ml) all receptors are occupied with ligand and are activated by the canonical dimerization mechanism. However, catalytic amplification can take place at sub-saturating EGF dose (20 ng/ml) because only a fraction of receptors will be occupied by ligand (20-30%; estimated by the ratio of EGF-Alexa647/mCitrine fluorescence at 20 ng/ml over 100 ng/ml EGF-Alexa647 per cell) (**Supplementary Figure 4a**). We therefore compared CONEGI and CONEGI-Y845F phosphorylation and conformational dynamics in cells that were stimulated with saturating or sub-saturating EGF dose. Upon stimulation with saturating EGF dose, CONEGI and CONEGI-Y845F exhibited similar rapid Y_1068_ phosphorylation reaching comparable levels (**Figure 4a; Supplementary Figure 4b**). CONEGI and CONEGI-Y845F also exhibited similar activation loop dynamics as apparent from the comparable change in τ over time (**Figure 4b; Supplementary Figure 4d**). This is consistent with the activation loop being rearranged to an open conformation in both CONEGI and its Y845F mutant by the canonical allosteric dimerization mechanism. A similar EGF-induced decrease in anisotropy for CONEGI and CONEGI-Y845F confirmed that Y845F mutation does not affect its dimerization (**Supplementary Figure 4c**).

**Figure 4:**
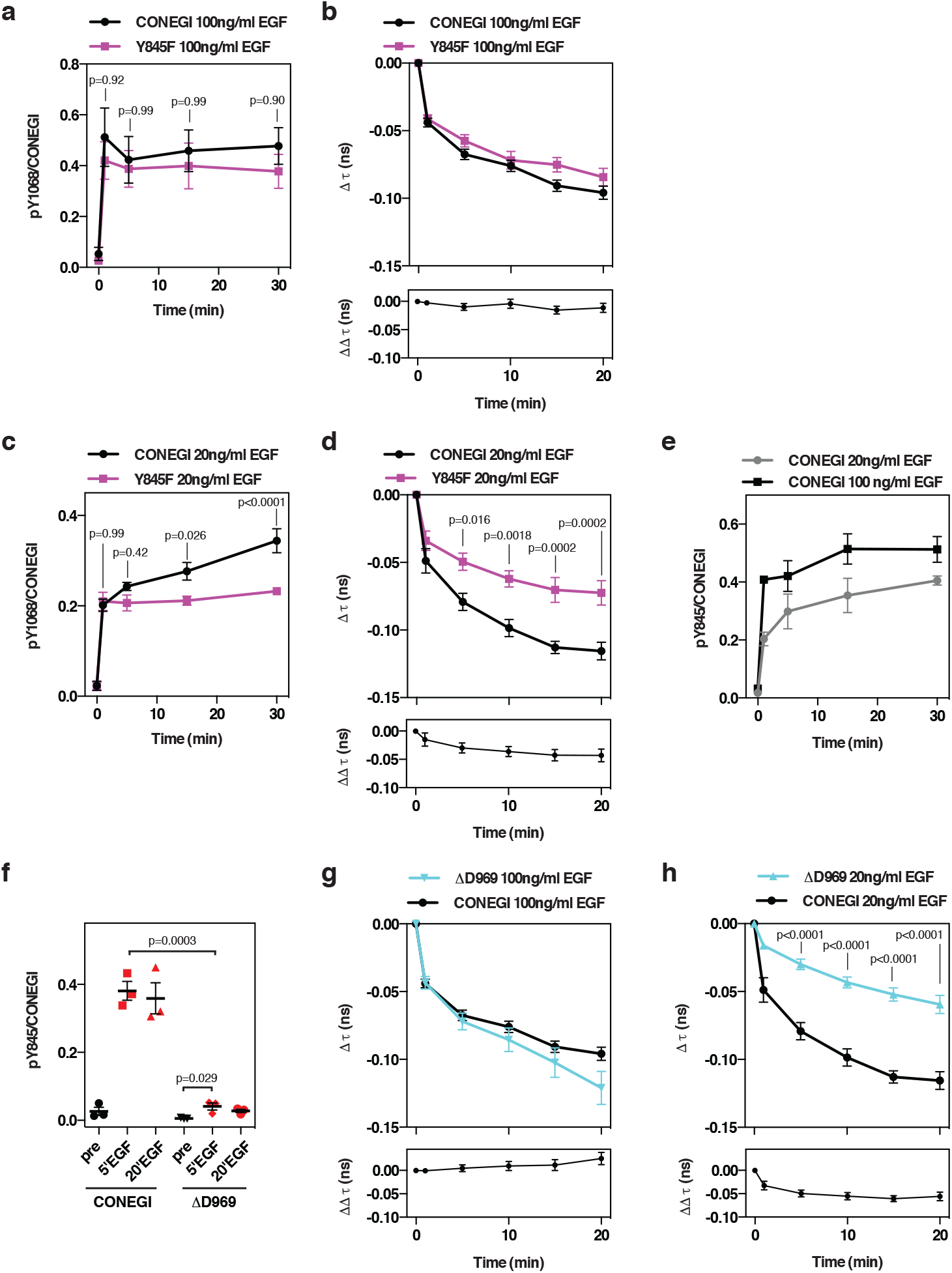
EGFR dimers can induce autocatalytic activation of EGFR monomers. (**a**) Relative Y_1068_ phosphorylation (pY_1068_/CONEGI) of CONEGI and CONEGI-Y845F upon stimulation with 100 ng/ml EGF by Western blot analysis (n=4) (**Supplementary Figure 4b**). (**b**) Change in Δτ of CONEGI and CONEGI-Y845F at the PM (upper panel) upon stimulation with 100 ng/ml EGF and the difference (ΔΔτ) between CONEGI and CONEGI-Y845F (lower panel) (CONEGI: n=19 cells; Y845F: n=8) (**Supplementary Figure 4d**). (**c**) Relative Y_1068_ phosphorylation (pY_1068_/CONEGI) of CONEGI and CONEGI-Y845F upon stimulation with 20 ng/ml EGF by Western blot analysis (n=4) (**Supplementary Figure 4b**). (**d**) Change in Δτ of CONEGI and CONEGI-Y845F at the PM upon stimulation with 20 ng/ml EGF (upper panel) and the difference (ΔΔτ) between CONEGI and CONEGI-Y845F (lower panel) (CONEGI: n=6 cells; Y845F: n=7) (**Supplementary Figure 4e**). (**e**) Relative Y_845_ phosphorylation (pY_845_/CONEGI) of CONEGI upon stimulation with 20 and 100 ng/ml EGF by Western blot analysis (n=3) (**Supplementary Figure 4f**). (**f**) Relative Y_845_ phosphorylation (pY_845_/CONEGI) of CONEGI and CONEGI-ΔD969 upon stimulation with 100 ng/ml EGF by Western blot analysis (n=3) (**Supplementary Figure 4h**). (**g,h**) Change in Δτ of CONEGI-ΔD969 and CONEGI at the PM upon stimulation with 20 (**h**) and 100 ng/ml EGF (**g**) (upper panel) and the difference (ΔΔτ) between CONEGI and CONEGI-ΔD969 (lower panel) (CONEGI: 20 ng/ml EGF: n=6 cells; 100 ng/ml EGF: n=19; ΔD969: 20 ng/ml EGF: n=9; 100 ng/ml: n=6) (**Supplementary Figure 4i**). Error bars: standard error of the mean. τ, fluorescence lifetime of mCitrine.

However, upon stimulation of cells with sub-saturating EGF dose, Y_1068_ phosphorylation of CONEGI reached within one minute ~39±9% of that at saturating EGF dose to further increase over time to ~72±12%, showing a clear amplification of Y_1068_ phosphorylation over time. This amplification was absent in CONEGI-Y845F, which after rapid initial activation remained stable at ~49±7% of that of CONEGI at saturating EGF (**Figure 4c; Supplementary Figure 4b**). Strikingly, the change in τ of CONEGI-Y845F upon sub-saturating EGF stimulation was significantly slower and of lesser magnitude as compared to that of CONEGI, which was comparable to that at saturating EGF (**Figure 4d; Supplementary Figure 4e**). This shows that EGF-activated CONEGI dimers induce an active conformation in unliganded CONEGI monomers by Y_845_ phosphorylation (**Figure 4e; Supplementary Figure 4f**). The question remained if Y_845_ is directly phosphorylated by the intrinsic kinase activity of EGFR or whether this happens indirectly by recruitment or activation of another tyrosine kinase that is dependent on C-terminal tyrosine auto-phosphorylation of EGFR^34^. To address this, we investigated Y_845_ phosphorylation and conformational dynamics of a C-terminal tail truncated CONEGI mutant (CONEGI-ΔD969). This mutant retained its ability to dimerize (**Supplementary Figure 4g**), but was severely impaired in Y_845_ phosphorylation (**Figure 4f, Supplementary Figure 4h**). At saturating EGF dose, CONEGI-ΔD969 followed closely the conformational dynamics of CONEGI (**Figure 4g; Supplementary Figure 4i**), consistent with activation by the allosteric dimerization mechanism. However, at sub-saturating EGF dose the change in τ of CONEGI-ΔD969 was significantly slower and of lesser magnitude as compared to that of CONEGI (**Figure 4h; Supplementary Figure 4i**). In fact, C-terminal truncation led to a similar attenuation of the conformational transition as the Y845F mutation (compare **Figure 4h** to **4d**). This shows that C-terminal auto-phosphorylation on EGFR is necessary to recruit or activate another tyrosine kinase that activates monomers by trans-phosphorylation on Y_845_ ^34^. These activated monomers can phosphorylate EGFR on the C-terminal tail, closing an autocatalytic loop.

## Discussion

Many insights in the ligand-induced allosteric dimer activation mechanism of EGFR were gained by structural studies that captured different conformations of the ECD and the TKD^2,5^. On the other hand, genetically encoded optical kinase substrate sensors provided information about spatially resolved EGFR phosphorylation dynamics^26,35^, but could not relate EGFR conformational dynamics to its phosphorylation activity. To bridge this gap, we developed the conformational sensor CONEGI that reports on conformational transitions in the activation loop of the EGFR TKD in living cells thereby enabling us to relate EGFR’s conformation to its phosphorylation activity and dimerization.

We designed multiple CONEGI variants in a hybrid approach using genetic code expansion in order to detect conformational dynamics in different key functional regions of the TKD. The limited availability of membrane-permeable dyes restricted the selection of an acceptor fluorophore with the spectral properties to form a FRET pair with mCitrine. This likely explains why existing conformational, FRET-based EGFR sensors are either based on protein labeling of the ECD^36,37^ or on *in vitro* labeling of intracellular domains^38^. The development of approaches to deliver membrane-impermeable dyes into cells^39^ or of novel membrane-permeable fluorophore species e.g. near-infrared silicon-rhodamine dyes^40^ will facilitate the generation of similar conformational FRET sensors in the future. Here, we proved BCNK-labeling with the membrane-permeable Atto590 to be stable and highly specific in living cells (**Figure 1d,e,g,h**). BCNK incorporation at the individual sites of the EGFR TKD affected kinase functionality to a different extent. Increased autonomous phosphorylation of CONEGI-712/713 and −713 might be caused by perturbation of inhibitory electrostatic interactions of kinase loop 2 with the PM^7^. The low EGF-induced fold-change in phosphorylation of CONEGI-737 could originate from hindrance of salt bridge formation between lysine 721 and glutamate 738 that is essential for adopting the active conformation^9^ or from perturbation of the αC-helix structure. In contrast, BCNK incorporation at position 851 in the activation loop was compatible with kinase function, probably because the activation loop is a highly flexible region and the incorporation site is sufficiently apart from the catalytically important DFG motif (830-832), Y_845_ and a β-strand that turns into a two-turn helix upon activation^10^.

With CONEGI we investigated if and by which mechanism catalytic EGFR activation occurs. The comparison in dimerization (by fluorescence anisotropy) and conformational transitions in the activation loop (by FLIM) of CONEGI with its Y845F mutant, not only showed the conformational transition driven by allosteric interactions in EGF-activated EGFR dimers^2^, but also that Y_845_ phosphorylation induces a conformational change in EGFR monomers. That this conformational transition induced by Y_845_ phosphorylation leads to a catalytically more active conformation was apparent from the increased Y_1068_ auto-phosphorylation on CONEGI monomers with respect to the CONEGI–Y845F mutant (**Figure 3e-i**). The PV-induced residual conformational transition in CONEGI-Y845F that correlated with a lower Y_1068_ phosphorylation suggests that besides Y_845_ phosphorylation, additional posttranslational reactions or interactions with other proteins could contribute to generate a catalytically competent conformation in EGFR monomers. At sub-saturating EGF dose, CONEGI Y_1068_ phosphorylation amplification correlated with the full conformational transition of its activation loop, whereas this amplification was absent in CONEGI-Y845F (**Figure 4c,d**). This showed that EGFR dimers can activate the tyrosine kinase activity of EGFR monomers by phosphorylation on Y_845_. The diminished Y_845_ phosphorylation and impaired conformational transition in C-terminal tail truncated CONEGI-ΔD969 monomers, further showed that these depend on auto-phosphorylation of the C-terminal tail (**Figure 4f,h**). EGFR autocatalysis can therefore occur by an Y_845_ phosphorylation induced catalytically active conformation in EGFR monomers that trans-phosphorylate EGFR monomers on their C-terminal tail. These auto-phosphorylation events on the C-terminal tail then induce the recruitment or activation of a tyrosine kinase that phosphorylates Y_845_ on EGFR monomers thereby generating a looped causality that amplifies EGFR phosphorylation. Src is known to be activated by C-terminally phosphorylated EGFR and capable of phosphorylating Y_845_ ^41,42^ and therefore likely involved in this autocatalytic amplification on EGFR.

Theoretically, the autocatalytic phosphorylation mechanism can generate only one collective state in which all receptors are fully active^43^. Therefore, the question arises how autocatalytic activation is regulated by interaction with other enzymatic activities. We previously demonstrated that continuous vesicular recycling of EGFR monomers through perinuclear areas with high protein tyrosine phosphatase (PTP) activity, such as that of the ER-associated PTP1B, counteracts spontaneous autocatalytic EGFR activation by Y_845_ dephosphorylation on recycling receptors^15^. The spatial separation of PTP activity from EGFR activity at the PM thereby provides a means via which cells can suppress autonomous autocatalytic activation while maintaining responsiveness to EGF. The collective response properties of EGFR at the PM however arise from the autocatalytic property of EGFR in conjunction with recursive interaction with PTPs in form of a double negative feedback^14,43,44^. Autocatalytic EGFR activation can lead to full EGFR phosphorylation at the PM above a threshold EGF concentration^14,45,46^. However, EGFR is organized in clusters of 100-200 nm in diameter with up to 250 EGFR monomers^33,47,48^ limiting the reach of autocatalysis. This could effectively create nanoscopic systems that can locally sense and robustly respond to growth factors.

## Online Methods

### Antibodies

#### Primary antibodies

Rabbit anti-EGFR (4267, Cell Signaling Technology, Danvers, MA), mouse anti-pY1068 (2236, Cell Signaling Technology, Danvers, MA), mouse anti-pY845 (558381, BD Biosciences, Heidelberg; Germany), mouse anti-phospho tyrosine (PY72) (P172.1, InVivo Biotech Services, Henningsdorf, Germany), mouse monoclonal anti-α-Tubulin (Sigma Aldrich, St. Louis, MO), rabbit anti-GAPDH (2118, Cell Signaling Technology, Danvers, MA), living colors rabbit anti-GFP (632593, Clontech, Mountain View, CA).

#### Secondary antibodies for Western blots

IRDye 680 donkey anti-mouse IgG (LI-COR Biosciences, Lincoln, NE), IRDye 800 donkey anti-rabbit IgG (LI-COR Biosciences, Lincoln, NE)

#### Secondary antibody for immunofluorescence

Alexa Fluor^®^ 647 donkey anti-mouse IgG (Thermo Fisher Scientific Inc., Waltham, MA)

### Plasmids

Restriction and ligation enzymes were purchased from New England Biolabs (NEB, Frankfurt am Main, Germany). mCitrine-N1, mCherry-N1 and mTurquoiseN1 were generated by insertion of AgeI/BsrgI PCR fragments of mCitrine, mCherry and mTurquoise cDNA in pEGFP-C1 (Clontech, Mountain View, CA). Fusions of EGFR and PTB domain with fluorescent proteins were generated through restriction-ligation of EGFR and PTB domain cDNA into the appropriate vector. To generate EGFR-QG-mCitrine mCitrine flanked with a linker sequence (LAAAYSSILSSNLSSDS-mCitrine-SDSSLNSSLISSYAAAL) was inserted between Q958 and G959 of EGFR. Site-directed mutagenesis PCR using *PfuUltra* High-Fidelity DNA polymerase (Agilent Technologies, Santa Clara, CA) replaced coding sequences in EGFR-QG-mCitrine with an amber codon (TAG) to generate BCNK incorporation sites and was also used to generate the CONEGI-Y845F mutant. NotI/NheI EGFR(TAG)-QG-mCitrine PCR fragments were inserted in a (U6-PylT*)_4_/EF1α plasmid previously described in ^23^. The BCNK incorporation system consists of two plasmids, (U6-PylT*)_4_/EF1α-PylRS to express the tRNA-synthetase and peRF1(E55D) for expression of a modified eukaryotic release factor 1 (eRF1). peRF1(E55D) was described earlier^23^ and (U6-PylT*)_4_/EF1α-PylRS was modified by adding a nuclear export sequence (NES) LALKLAGLDIGS attached via a flexible 4x(SGGGGS) linker to the N-terminus of PylRS. The plasmid was created from a previously reported construct^49^ by inverse PCR followed by insertion of the 4x(SGGGGS) linker via blunt end ligation. All constructs were sequence verified and tested for correct expression.

### Reagents

Human EGF (Peprotech, Hamburg, Germany) was shock frozen at a concentration of 100 μg/ml in PBS + 0.1% BSA and stored at −80°C. Pervanadate was freshly prepared by adding sodium orthovanadate (S6508, Sigma Aldrich, St. Louis, MO) to H_2_0_2_ (30%) according to ^30^. BCNK was described earlier in ^24^. Site-specific C-terminal labeling of hEGF with Alexa647-maleimide (Life Technologies, Darmstadt, Germany) was carried out as described previously in ^50^. Tet-Atto590 was synthesized by conjugating Atto590-NHS ester (79636, Sigma Aldrich, St. Louis, MO) to tet-NH_2_ in the presence of N,N-Diisopropylethylamine (DIPEA). The product was purified using HPLC and the identity of the compound confirmed by mass spectroscopy. Lapatinib (231277-92-2, Cayman Chemical, Ann Arbor, MI) was solubilized in Ethanol to a stock concentration of 10 mM and stored at 4°C.

### Crystal structure

PDB files of crystal structures of the active and inactive EGFR TKD were downloaded from the RSCB protein data bank. Alignment of the crystal structures and measurement of the distances between insertion sites were performed using the program MacPyMOL (http://www.pymol.org).

### Cell culture, transfections and site-specific labeling of EGFR

Identity of HEK293T cells (ATCC CRL-11268) was determined by DNA profiling using 8 different and highly polymorphic short tandem repeat (STR) loci and testing for the presence of mitochondrial DNA sequences from rodent cells as mouse, rat, chinese and syrian hamster (DSMZ) and tested regularly for mycoplasma contamination using MycoAlert Mycoplasma detection kit (Lonza). HEK 293T cells were grown in Dulbecco’s Modified Eagle’s Medium (DMEM) supplemented with 10% fetal bovine serum (FBS), 2 mM L-glutamine and 1% non-essential amino acids (NEAA) and maintained at 37**°**C in 5% CO_2_. Transfection of mammalian cells with plasmid DNA was achieved using Fugene^®^6 according to the manufacturers protocol. HEK293T cells in one well of an 8-well Labtek (for confocal microscopy and FLIM) were transfected with 250 ng of the EGFR expression plasmid, 100 ng (U6-PylT*)_4_/EF1α-PylRS and 100 ng peRF1(E55D). Cells in one well of a 24-well dish (for Western blot analysis) were transfected with 500 ng of the EGFR expression plasmid, 100 ng (U6-PylT*)_4_/EF1α-PylRS and 100 ng peRF1(E55D). HEK293T cells in MatTeks (for anisotropy) were transfected with 1000 ng of the EGFR expression plasmid, 100 ng (U6-PylT*)_4_/EF1α-PylRS and 100 ng peRF1(E55D). Media was replaced by fresh media supplemented with 1 mM BCNK before addition of the transfection mix to cells. Cells were incubated overnight at 37°C with 5% CO_2_ and BCNK was washed out on the next day. Labeling was performed using 400 nM tet-Atto590 in DMEM for 20 min at room temperature (RT). The washout of free diffusing tet-Atto590 was carried out for at least 2 h with media exchange every 15-20 min.

### Western blotting and in-gel fluorescence

Cells were lysed in ready-made Cell Lysis Buffer (9803, Cell Signaling Technology, Danvers, MA) supplemented with Complete Mini EDTA-free protease inhibitor (Roche Applied Science, Heidelberg, Germany) and 100 μl phosphatase inhibitor cocktail 2 and 3 (P5726 and P0044, Sigma Aldrich, St.Louis, MO). Following lysis, samples were cleared by centrifugation for 10 min, 13,000 rpm at 4°C. Bis-Tris-PAGE was performed using the X-cell II mini electrophoresis apparatus (Life Technologies, Darmstadt, Germany) according to the manufacturers instructions. Proteins were transferred to preactivated polyvinylidene difluoride (PVDF) membranes (Merck Chemicals, Darmstadt, Germany) and incubated with the respective primary antibodies at 4**°**C overnight. Detection was performed using species-specific IR-Dye 800 CW and IR-Dye 680 secondary antibodies (LI-COR Biosciences, Bad Homburg vor der Höhe, Germany) and the Odyssey Infrared Imaging System (LI-COR Biosciences, Bad Homburg vor der Höhe, Germany). Eventually, Bis-Tris gels were imaged with the Typhoon Trio Variable Mode Imager (GE Healthcare, Buckinghamshire, UK) to detect fluorescently labeled proteins before Western blotting. Atto590 was excited using a 532 nm laser and fluorescence emission was detected with a 610/30 BP filter at a resolution of 100 microns.

### Immunofluorescence

Cells were fixed with 4% paraformaldehyde (Roti^®^-Histofix 4%, Carl Roth GmbH, Karlsruhe, Germany) for 10 min at RT and permeabilized for 5 min with 0.1% Triton X-100 in TBS. Background staining was blocked by incubation with Odyssey^®^ Blocking Buffer (LI-COR Biosciences, Lincoln, NE) for 1 h at 4°C. Primary antibodies diluted in Odyssey^®^ Blocking Buffer were applied overnight at 4°C and secondary antibodies for 1 h at RT. Fixed cells were imaged in TBS at 37°C. In background-subtracted images, masks for the PM of single cells were generated and the mean fluorescence intensity for each channel was measured in ImageJ (http://imagej.nih.gov/ij/). The relative phosphorylation on Y_1068_ was determined per cell and the mean values of Y_1068_/EGFR-QG-mCitrine variants were calculated.

### Fluorescence microscopy

#### Olympus FV1000

Confocal images at the Olympus FV1000 equipped with a 60x/1.35 NA Oil UPLSApo objective (Olympus, Hamburg, Germany) and a temperature controlled incubation chamber (EMBL, Heidelberg, Germany) were acquired in sequential mode frame by frame with 2x line averaging. The pinhole was set to 2.5 airy units. mCitrine was excited with a 488 nm Argon laser (GLG 3135, Showa Optronics, Tokyo, Japan), mCherry/Atto590 with a 561 nm DPPS laser (85-YCA-020-230, Melles Griot, Bensheim, Germany) and Alexa647 with a 633 HeNe laser (05LHP-991, Melles Griot, Bensheim, Germany) using a DM405/488/561/633 dichroic mirror. mCitrine fluorescence was detected between 498-551 nm using the SDM560 beam splitter. Atto590 fluorescence was detected in the bandwidth of 575-675 nm and Alexa647 fluorescence between 643-743 nm. Live cells were imaged in imaging media at 37°C and 5% CO_2_ and stimulated with 20 ng/ml EGF, 100 ng/ml EGF or 0.33 mM PV.

### Fluorescence lifetime imaging microscopy

FLIM data were obtained with the Olympus FV1000 laser scanning microscope (Olympus, Hamburg, Germany) equipped with an external unit, PicoQuant’s compact FLIM and FCS upgrade kit laser scanning microscopes (Picoquant GmbH, Berlin, Germany) using a 60x/1.35 NA Oil UPLSApo objective (Olympus, Hamburg, Germany). Pulsed lasers were coupled to the FV1000 through an independent port and controlled with SepiaII software (Picoquant GmbH, Berlin, Germany). Detection of photons was achieved using a single photon avalanche diode (PDM Series, MPD, Picoquant GmbH, Berlin, Germany) and timed using a single photon counting module (PicoHarp 300, Picoquant GmbH, Berlin, Germany). Using the SymPhoTime software V5.13 (Picoquant GmbH, Berlin, Germany) images were collected with an integration time of ~2 min collecting ~3.0-5.0 × 10^6^ photons. mCitrine was excited by a 507 nm pulsed laser (at 67% of maximum laser power) (LDH 507, Picoquant GmbH, Berlin, Germany) and fluorescence of mCitrine was collected using a narrow-band emission filter (HQ 537/26, Chroma, Olching, Germany). FLIM data were analysed using the global analysis code described in ^51^ to obtain images of the mean fluorescence lifetime τ. Pixels with a total number of counts less than 50 photons were excluded from analysis. Masks for single cells or for the PM of single cells were generated and the mean fluorescence τ was measured in ImageJ (http://imagej.nih.gov/ij/).

### Widefield anisotropy

Anisotropy microscopy was performed on an Olympus IX81 inverted microscope (Olympus, Hamburg, Germany) equipped with a 20x/0.7 NA air objective using an Orca CCD camera (Hamamatsu Photonics, Hamamatsu City, Japan) and an incubation chamber (EMBL, Heidelberg, Germany). mCitrine and Atto590 were excited using a MT20 illumination system. A linear dichroic polarizer (Meadowlark optics, Frederick, CO) was implemented in the illumination path of the microscope, and two identical polarizers were placed in an external filter wheel at orientations parallel and perpendicular to the polarization of the excitation light. For each field of view two images were taken, one with the emission polarizer oriented parallel to the excitation polarizer (*I*_∥_) and one with the emission polarizer oriented perpendicular to the excitation polarizer (*I*_⊥_). The fluorescence anisotropy (r^i^) in each pixel *i* was calculated according to:

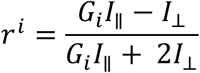

To determine the G-factor (G_i_) parallel and perpendicular images of the fluorophore fluorescein were taken in solution. Fluorescein’s anisotropy is close to zero and therefore allows calculating G_i_ by building the ratio of the perpendicular over the parallel intensities. The CellR software supplied by the microscope manufacturer (Olympus, Hamburg, Germany) controlled data acquisition. Live cells were imaged in vitamin-free media at 37°C and 5% CO_2_ and stimulated with either 100 ng/ml EGF or 0.33 mM pervanadate.

## Statistical analysis

All results are expressed as mean ± S.E.M or in Tukey box plots. Statistical analysis was performed with GraphPad Prism, version 6.0e for Mac (GraphPad Software, La Jolla, CA, USA). Statistical significance was estimated either by unpaired two-tailed t tests or by two-way analysis of variance (ANOVA).

## Data availability

The authors declare that data supporting the findings of this study are included within the paper and its supplementary information files or are available from the corresponding author upon reasonable request.

## Acknowledgement

This project was partially funded by the following grants: EMBO Short-Term Fellowship (ASTF No: 122-2015) to MB, MC_U105181009 and MC_UP_A024_1008 to JWC and MRC-Nikon Case Studentship to VB. We thank Dr A Sachdeva for synthesizing tetrazine-Atto590, Dr P Bieling and Dr A Krämer for critically reading the manuscript.

## Author contributions

PIHB conceived the project. MB designed the conformational sensor, cloned sensor constructs and acquired and analyzed the data. CU and VB provided previously unpublished expression constructs and instruction on site-specific fluorophore labeling. MG analyzed anisotropy data. MB, JWC and PIHB wrote the manuscript with input from all authors.

## Competing financial interests

The authors declare no competing financial interests.

## Supplementary information

**Supplementary Figure 1:**
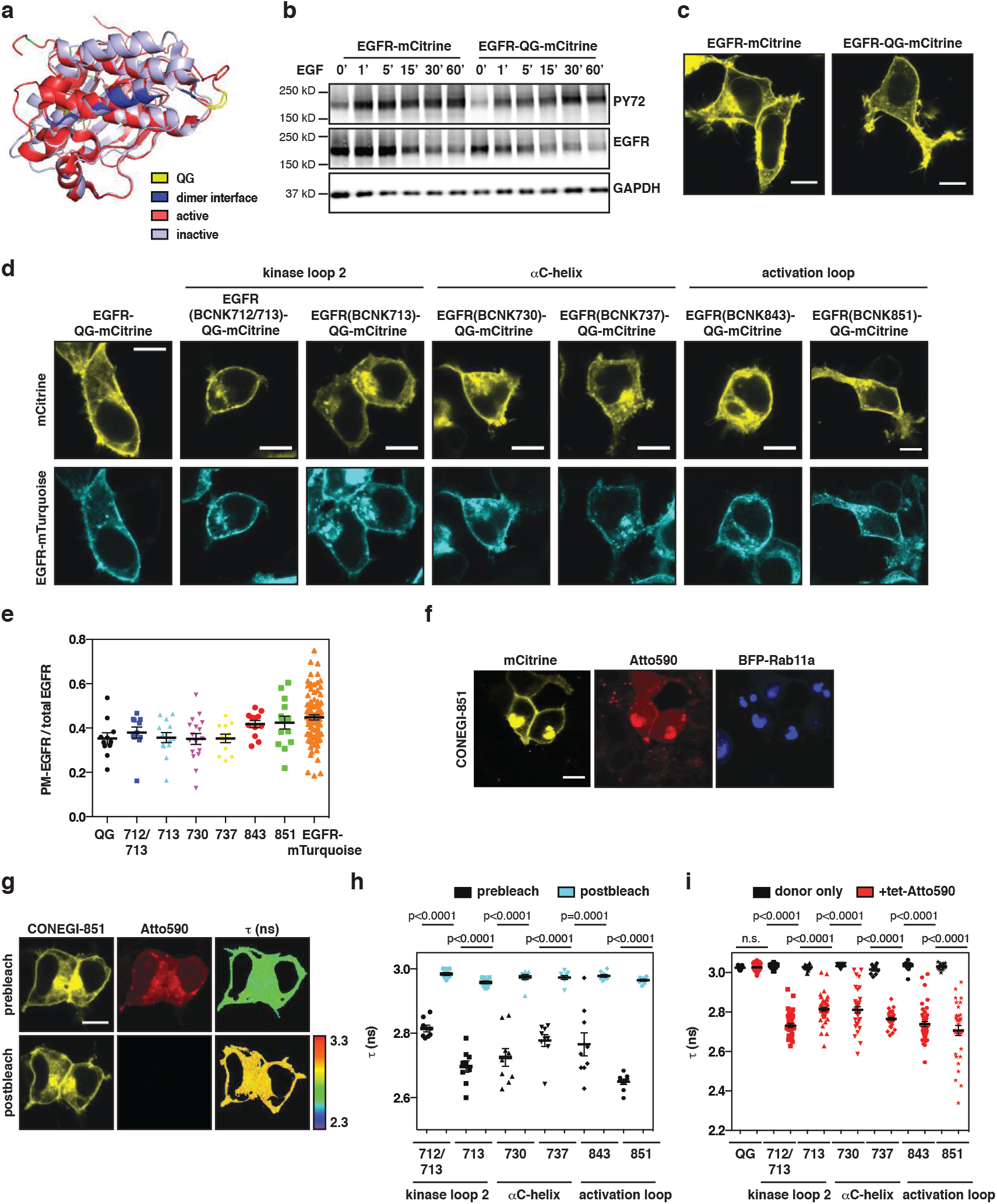
Design and characterization of CONEGI. (**a**) Alignment of an active (red; PDB: 2J5F) and inactive (cyan; PDB: 2GS7) crystal structure of the EGFR TKD. mCitrine insertion site (yellow) and dimerization interface (blue) are indicated. (**b**) Phosphorylation of EGFR-mCitrine and EGFR-QG-mCitrine in HEK293T cells upon EGF stimulation. Western blots using whole cell lysates were probed with anti-EGFR, anti-PY72 and anti-GAPDH (**Figure 1c**). (**c**) Representative fluorescence images of EGFR-mCitrine and EGFR-QG-mCitrine. (**d**) Representative fluorescence images of HEK293T cells co-expressing EGFR-mTurquoise and EGFR-QG-mCitrine or EGFR(BCNKXXX)-QG-mCitrine variants. (**e**) Quantification of the relative PM-EGFR fraction (PM-EGFR/total EGFR) of EGFR-QG-mCitrine (n=11 cells), EGFR(BCNKXXX)-QG-mCitrine variants (712/713: n=11; 713: n=13; 730: n=17; 737: n=12; 843: n=12; 851: n=14) and EGFR-mTurquoise (n=90). (**f**) Representative fluorescence images of HEK293T cells co-expressing CONEGI-851 (mCitrine and Atto590 fluorescence) and BFP-Rab11a. (**g**) Representative mCitrine and Atto590 fluorescence images of CONEGI-851 before and after photobleaching Atto590 and corresponding τ images. (**h**) Mean τ in CONEGIs (712/713: n=8 cells; 713: n=9; 730: n=9; 737: n=9; 843: n=9; 851: n=9) at the PM before and after photobleaching Atto590. (**i**) Mean τ of EGFR-QG-mCitrine or EGFR(BCNKXXX)-QG-mCitrine variants (QG: n=11 cells; 712/713: n=13; 713: n=12; 730: n=12; 737: n=9; 843: n=12; 851: n=14) and their corresponding CONEGI variants (QG: n=42; 712/713: n=36; 713: n=43; 730: n=36; 737: n=30; 843: n=44; 851: n=32) at the PM. Scale bars: 10 μm. EGF stimulation, 100 ng/ml. Error bars: standard error of the mean. τ, fluorescence lifetime of mCitrine.

**Supplementary Figure 2:**
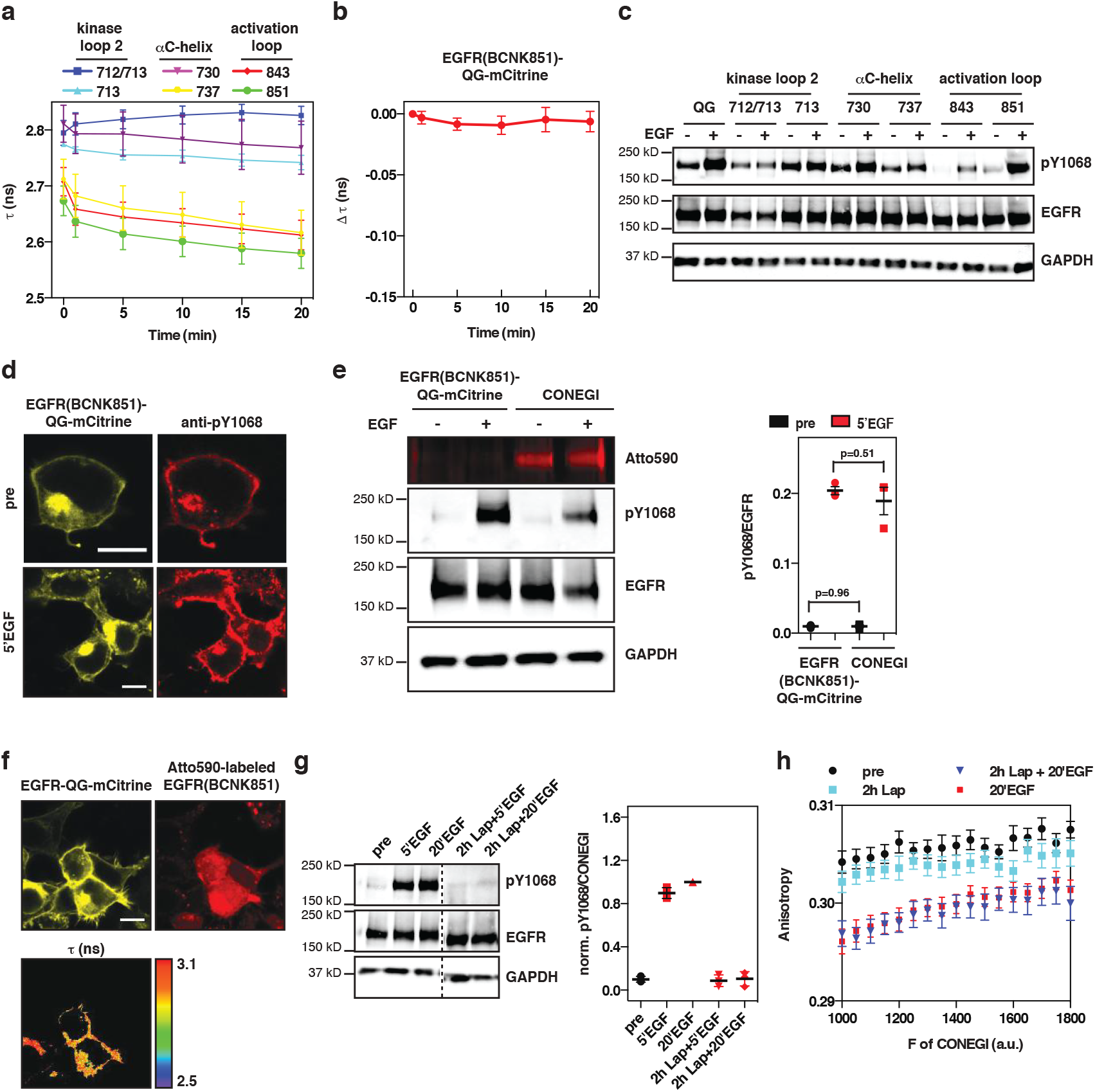
CONEGI reports on conformational transitions of the activation loop. (**a**) Change in mean τ of CONEGI variants at the PM upon EGF stimulation (QG: n=5 cells; 712/713: n=7; 713: n=7; 730: n=7; 737: n=7; 843: n=6; 851: n=31). (**b**) Change in Δτ of EGFR(BCNK851)-QG-mCitrine at the PM upon EGF stimulation (n=3 cells). (**c)** Representative Western blot showing Y_1068_ phosphorylation of EGFR-QG-mCitrine and CONEGIs upon 5 min EGF stimulation. Blots were probed with anti-pY_1068_, anti-EGFR and anti-GAPDH (**Figure 2e**). (**d**) Representative fluorescence images of mCitrine and anti-pY_1068_. HEK293T cells expressing EGFR(BCNK851)-QG-mCitrine were immunostained against pY_1068_ upon EGF stimulation (**Figure 2g**). (**e**) Representative fluorescence image and Western blot of HEK293T cells expressing EGFR(BCNK851)-QG-mCitrine left unlabeled or 20 min labeled with tet-Atto590 show Atto590 fluorescence, Y_1068_ phosphorylation and EGFR expression upon EGF stimulation. Blots were probed with anti-pY_1068_, anti-EGFR and anti-GAPDH (left). Corresponding quantification of Y_1068_ phosphorylation (pY_1068_/EGFR) of EGFR(BCNK851)-QG-mCitrine and CONEGI (n=3) (right). (**f**) Representative fluorescence images of HEK293T cells co-expressing EGFR-QG-mCitrine and Atto590-labeled EGFR(BCNK851) and corresponding τ (**Figure 2h**). (**g**) Representative Western blot showing Y_1068_ phosphorylation of HEK293T cells expressing CONEGI upon EGF stimulation in absence or presence of 1 μM Lapatinib (Lap). Blots were probed with anti-pY_1068_, anti-EGFR and anti-GAPDH (left). Corresponding quantification of relative Y_1068_ phosphorylation (pY_1068_/CONEGI) of CONEGI (right) (n=3) (**h**) mCitrine fluorescence anisotropy of CONEGI versus binned mean fluorescence intensity (F of CONEGI) per pixel upon EGF stimulation in presence or absence of 1 μM Lap (N=3 experiments) (**Figure 2j**). Scale bars: 10 μm. EGF stimulation, 100 ng/ml. Error bars: standard error of the mean. τ, fluorescence lifetime of mCitrine.

**Supplementary Figure 3:**
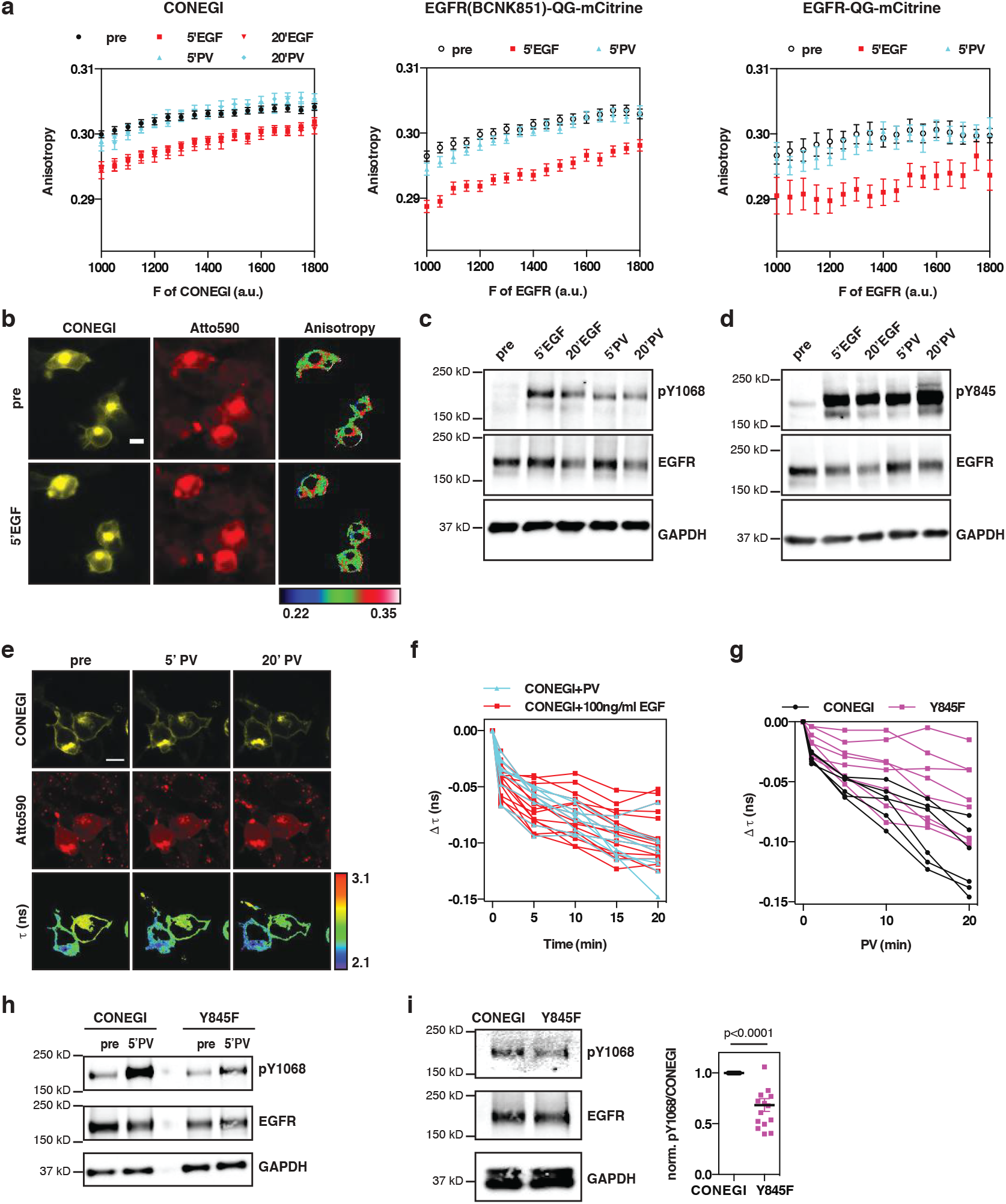
EGFR monomers phosphorylated on Y845 adopt an active conformation. (**a**) mCitrine fluorescence anisotropy of CONEGI (left), EGFR(BCNK851)-QG-mCitrine (middle) and EGFR-QG-mCitrine (right) versus its binned mean fluorescence intensity (F of CONEGI or EGFR-QG-mCitrine variants) per pixel in HEK293T cells upon EGF or pervanadate (PV) treatment (N=3 experiments/variant) (**Figure 3a**). (**b**) Representative mCitrine and Atto590 fluorescence images of CONEGI and corresponding mCitrine fluorescence anisotropy images upon EGF stimulation. (**c,d**) Representative Western blots of HEK293T lysates transfected with CONEGI showing Y_1068_ (**c**) or Y_845_ (**d**) phosphorylation and expression of CONEGI (EGFR) upon EGF or PV treatment. Blots were probed with anti-pY_1068_ or anti-pY_845_, anti-EGFR and anti-GAPDH (**Figure 3b,c**). (**e**) Representative mCitrine and Atto590 fluorescence images and corresponding τ of CONEGI upon PV treatment. (**f**) Change in Δτ of CONEGI at the PM upon EGF or PV treatment in individual cells (**Figure 3d**). (**g**) Change in Δτ of CONEGI and CONEGI-Y845F in individual cells upon PV treatment (**Figure 3f**). (**h**) Representative Western blot on HEK293T lysates transfected with CONEGI or CONEGI-Y845F upon PV treatment show Y_1068_ phosphorylation and EGFR expression. Blots were probed with anti-pY_1068_, anti-EGFR and anti-GAPDH (**Figure 3e**). (**i**) Representative Western blot (left) on HEK293T lysates expressing CONEGI or CONEGI-Y845F show Y_1068_ phosphorylation and their expression (EGFR). Blots were probed with anti-pY_1068_, anti-EGFR and anti-GAPDH. Corresponding normalized Y_1068_ phosphorylation (pY_1068_/CONEGI) of CONEGI and CONEGI-Y845F in absence of ligand (right; n=15). Scale bars: 10 μm. EGF stimulation, 100 ng/ml. PV treatment, 0.33 mM. Error bars: standard error of the mean. τ, fluorescence lifetime of mCitrine.

**Supplementary Figure 4:**
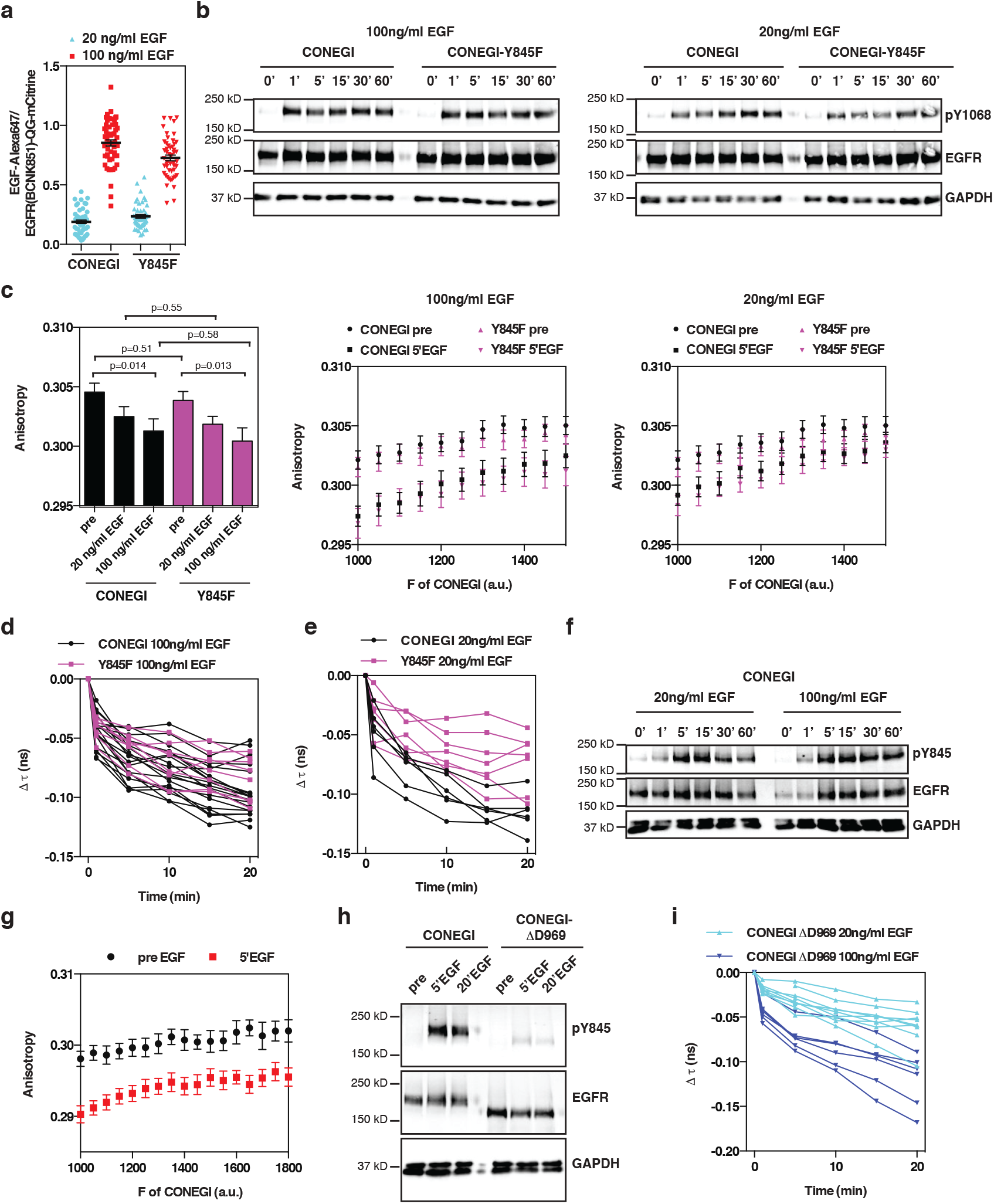
EGFR dimers can activate autocatalytic activation on EGFR monomers. (**a**) Relative ligand occupancy (EGF-Alexa647/EGFR(BCNK851)-QG-mCitrine) of EGFR-(BCNK851)-QG-mCitrine (20 ng/ml EGF: n=52 cells; 100 ng/ml EGF: n=53) and its Y845F mutant (20 ng/ml EGF: n=50; 100 ng/ml EGF: n=50) at the PM upon stimulation with 20 or 100 ng/ml EGF-Alexa647 . (**b**) Representative Western blots on HEK293T lysates expressing CONEGI or CONEGI-Y845F upon stimulation with 100 (left) or 20 ng/ml EGF (right). Blots were probed with anti-pY_1068_, anti-EGFR and anti-GAPDH (**Figure 4a,c**). (**c**) Mean mCitrine fluorescence anisotropy of CONEGI and CONEGI-Y845F upon 5 min stimulation with 20 or 100 ng/ml EGF (left). mCitrine fluorescence anisotropy of CONEGI or CONEGI-Y845F versus its binned mean fluorescence intensity (F of CONEGI) per pixel in HEK293T cells before and after 5 min stimulation with 100 (middle) or 20 ng/ml EGF (right) (N=3 experiments). (**d,e**) Change in Δτ of CONEGI or CONEGI-Y845F at the PM in individual HEK293T cells upon stimulation with 100 (**d**) or 20 ng/ml EGF (**e**) (**Figure 4b,d**). (**f**) Representative Western blot on HEK293T lysates expressing CONEGI upon stimulation with 20 or 100 ng/ml EGF. Blots were probed with anti-pY_845_, anti-EGFR and anti-GAPDH (**Figure 4e**). (**g**) mCitrine fluorescence anisotropy of CONEGI-ΔD969 versus its binned mean fluorescence intensity (F of CONEGI) per pixel upon stimulation with 100 ng/ml EGF (N=3 experiments). (**h**) Representative Western blot on HEK293T lysates expressing CONEGI or CONEGI-ΔD969 upon stimulation with 100 ng/ml EGF. Blots were probed with anti-pY_845_, anti-GFP (EGFR) and anti-GAPDH (**Figure 4f**). (**i**) Change in Δτ of CONEGI-ΔD969 at the PM in individual HEK293T cells upon stimulation with 20 or 100 ng/ml EGF (**Figure 4g,h**). Error bars: standard error of the mean.

**Supplementary Table 1:** FRET efficiencies of CONEGI variants. Estimated distances (including linker length) between mCitrine insertion site and each BNCK incorporation site as well as estimated and experimentally obtained FRET efficiencies for each COENGI variant. To estimate FRET efficiencies the orientation factor κ^2^ and the refractive index were assumed to be 2/3 and 1.4.

| BCNK incorporation site | Distance (Å) | Estimated FRET efficiency | Experimentally obtained FRET efficiency |
| --- | --- | --- | --- |
| 712/713 | 80.1 | 0.14 | 0.108 ± 0.006 |
| 713 | 80.1 | 0.14 | 0.112 ± 0.011 |
| 730 | 85.2 | 0.1 | 0.105 ± 0.008 |
| 737 | 79.7 | 0.14 | 0.105 ± 0.007 |
| 843 | 81.9 | 0.12 | 0.122 ± 0.006 |
| 851 | 73.7 | 0.21 | 0.143 ± 0.008 |

